# Iota oscillations (25-35 Hz) during wake and REM sleep in children and young adults

**DOI:** 10.1101/2024.08.06.606898

**Authors:** Sophia Snipes

## Abstract

High-frequency brain oscillations in humans are currently categorized into beta (13-30 Hz) and gamma (>30 Hz). Here, I introduce a new class of oscillations between 25 and 35 Hz, which I propose to call “iota.” Iota oscillations have low amplitudes but can still be measured with surface electroencephalography (EEG). Within an individual, iota has a narrow spectral bandwidth of 2-4 Hz, thus distinguishing it from broadband beta and gamma. Iota oscillations occur as sustained bursts during both wakefulness and REM sleep but do not appear during NREM sleep. They are only found in a minority of individuals, more in children than in adults. Overall, iota oscillations are challenging to detect but could serve as a marker of both brain development and states of vigilance.

## 2 INTRODUCTION

Oscillations are a fundamental aspect of human EEG activity and are categorized by their frequency: *delta* (0.5-4 Hz), *theta* (4-8 Hz), *alpha* (8-13 Hz), *sigma* (11-16 Hz), *beta* (13-30 Hz) and *gamma* (30-80 Hz).^1,2^ In addition to frequency, oscillations are characterized by their location in the brain, their waveform, and the cognitive or conscious states during which they appear. For example, alpha activity occurs primarily during relaxed wakefulness with eyes closed, it is measured in occipital channels, and it appears as long trains of near-sinusoidal oscillations.^3^ Sigma activity occupies an overlapping band but is only found during non-rapid-eye-movement (NREM) sleep, it appears in 0.5-2 s bursts known as “spindles”, and is measured primarily in central and frontal channels.^2–4^

Here, I describe a new oscillation, *iota*, which is likewise defined by a characteristic frequency, location, waveform, and conscious states. Iota oscillations are between 25 and 35 Hz, thus at the intersection between beta and gamma. This paper will demonstrate how iota differs from these neighboring oscillations, while also excluding potential artefacts. Finally, I will discuss prior literature that may relate specifically to iota and examine why these oscillations may have otherwise gone unreported until now.

To define iota, this paper uses freely available wake EEG data from diverse pediatric patient populations,^5^ and the open-source *fitting oscillations & one over f* (FOOOF) algorithm.^6^ To identify the conscious states in which iota occurs, sleep data from young healthy adults was analyzed.^7^

## 3 METHODS

### 3.1 Data

#### 3.1.1 Wake data from New York children

The Healthy Brain Network (HBN) dataset from the Child Mind Institute (https://data.healthybrainnetwork.org/) was collected in New York, USA, using a community-referred recruitment model, aimed at families who had concerns about psychiatric symptoms of their children. High-density EEG was recorded in a sound-shielded room at 500 Hz using 128-channel sponge-based Geodesic HydroCel nets from Electrical Geodesics Inc. (EGI). Recordings were referenced to Cz, impedances kept below 40 kOhm. Each recording lasted 5 minutes, with participants opening and closing their eyes at various timepoints.

3321 EEG recordings were obtained from releases 1-11. After preprocessing, the final dataset comprised 2496 individuals, 36% female, 13% left-handed, mean age of 10.4 (5-22). The diagnostic categories included: neurodevelopmental disorders (62%); no diagnosis indicated (14.2%); anxiety disorders (13.6%); depressive disorders (3.6%), disruptive, impulse control and conduct disorders (2.1%); and trauma and stressor related disorders (1.3%). All other categories each comprised less than 1% of the dataset. Neuro-developmental disorders included primarily attention-deficit hyperactivity disorder (ADHD; 69.9%), learning disorder with impairment in reading (10.9%), autism spectrum disorder (10.5%), and language disorder (2.9%). Further details of the dataset are provided in Alexander et al., (2017).^5^

#### 3.1.2 Sleep data from Zurich adults

Baseline sleep EEG data was collected from young healthy adults in Zurich, Switzerland, as part of a sleep deprivation study. The dataset comprised 19 participants, screened for good physical and mental health, good sleep quality, regular circadian rhythms, and some degree of vulnerability to sleep deprivation. Participants were between 18 and 26 years old (mean=23), 10 female, three left-handed. Further details are provided in Snipes et al., (2022, 2024).^7,8^

High-density EEG was recorded with 128-channel gel-based EGI Geodesic HydroCel nets, connected to DC BrainAmp Amplifiers and recorded with Brainvision Recorder software. Data was recorded at 1000 Hz sampling rate and Cz reference. Impedances were kept below 25 kOhm at the beginning of the night.

### 3.2 Analysis

All preprocessing and analyses were performed in MATLAB 2023b, using the EEGLAB toolbox,^9^ the FOOOF toolbox,^6^ and custom scripts (https://github.com/snipeso/iota-preprint, https://github.com/snipeso/eeg-oscillations).

#### 3.2.1 Wake

Wake data was preprocessed with the same fully automatic procedure described in Snipes et al., (2024).^10^ Very briefly, data was filtered between 0.5 and 50 Hz, further notch filtered at 60, 120, and 180 Hz to remove line noise and harmonics, and downsampled to 250 Hz. Timepoints or channels containing major movement or unphysiological artifacts were removed, data was re-referenced to the average, and independent component analysis (ICA) was used to remove eye, muscle, and heartbeat artifacts, using EEGLAB’s *iclabel* function to automatically classify components.

Spectral power was calculated for each channel in 20 s epochs using the *pwelch* function, with 4 s Hanning windows, 50% overlap. Data was smoothed over 2 Hz using the *lowess* method. As a final step to remove artefacts, FOOOF was applied to the power spectra of each channel and each epoch. This channel/epoch was removed if certain thresholds weren’t met. The aperiodic signal was fitted between 3-50 Hz. 3 Hz avoided lingering eye artifacts, and above 50 Hz the aperiodic slope changed, becoming less steep (see Figure 6D). For this reason, 50 Hz was also the cutoff for the low-pass filter. Otherwise, default settings were used (peak width: [0.5 12], max number of peaks: inf; minimum peak height: 0; peak threshold: 2; aperiodic mode: fixed). Datapoints were excluded if the model’s mean absolute error was >0.1 or r-squared <0.98. Additionally, epochs from a given channel were excluded if aperiodic exponents were outside values of 0-3.5, or offsets outside 0-4, which are the physiological ranges for wake data in this age group.^10^ If there were fewer than 80 artifact-free channels in an epoch, then the whole epoch was excluded.

After all artefacts were removed, power spectra were averaged over channels then epochs. FOOOF was then applied with the same parameters as before, to extract periodic power and periodic peaks for each recording. Periodic power was calculated by subtracting the aperiodic signal from the log-power signal (Figure 1). Periodic peaks were provided directly by the FOOOF algorithm; each peak was defined by its center frequency, its amplitude, and its bandwidth. Peaks with the outer edge of the bandwidth extending beyond the 3-50 Hz range were excluded from further analysis, as these typically reflected fitting imperfections. More details on analyses are provided in the figure legends.

**Figure 1:**
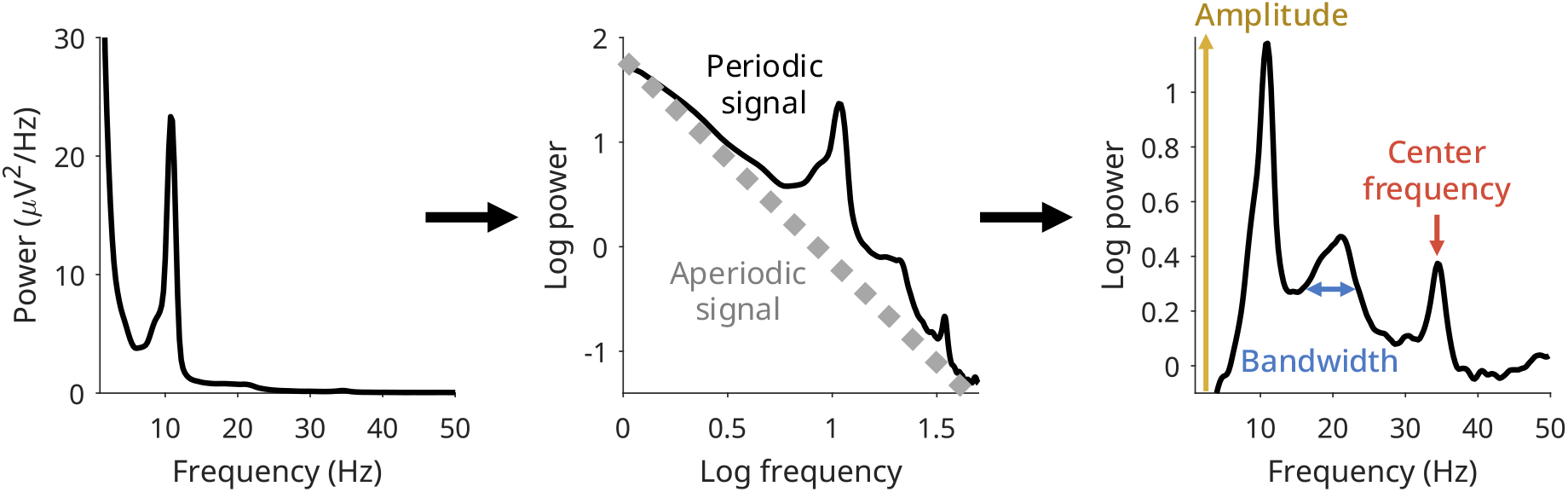
Detection of periodic peaks using the FOOOF algorithm. The EEG power spectrum (left) is transformed on a log-log scale, and a line is fitted to capture the background aperiodic activity. This is subtracted, and gaussians are fit iteratively until all of the “periodic power” is accounted for.^6^ Each gaussian is characterized by a center frequency, a bandwidth (2*std of the gaussian), and its amplitude. The example above includes both narrowband alpha and iota peaks, and a broadband beta peak. This recording produced three points in Figure 2B.

#### 3.2.2 Sleep

Sleep was scored manually using standard American Academy of Sleep Medicine (AASM) scoring criteria on 20 s epochs.^3^ Scores included wake, REM sleep, and three NREM sleep stages, N1, N2, and N3. The sleep EEG data was cleaned by first mean-centering each channel, then the data was low-pass filtered at 100 Hz using EEGLAB’s *pop_eegfiltnew* function, downsampled to 250 Hz, and high-pass filtered at 0.2 Hz (Kaiser method; stop-band frequency = 0.1; stopband attenuation = 60; passband ripple = 0.05). A FIR Kaiser filter was further used to remove 50 Hz line noise and harmonics. Epochs and channels containing artifacts were removed using a semi-automated approach.^11^ A channel or epoch was completely removed if more than 10% of the signal was marked as an artefact. Missing channels were interpolated (EEGLAB’s *pop_interp* function), then the EEG was re-referenced to the average.

The sleep data was analyzed first using the same procedure as for the wake EEG, averaging spectral power separately for each sleep stage. The investigated frequency range was shortened to 3-45 Hz to avoid the European 50 Hz line noise. In a second analysis, FOOOF was applied to each channel and epoch individually (Figure 7).

## 4 RESULTS

### 4.1 Wake iota

#### 4.1.1 Prevalence

Iota activity was detected in the power spectra of resting wake EEG data of the HBN pediatric dataset (Figure 2).

**Figure 2:**
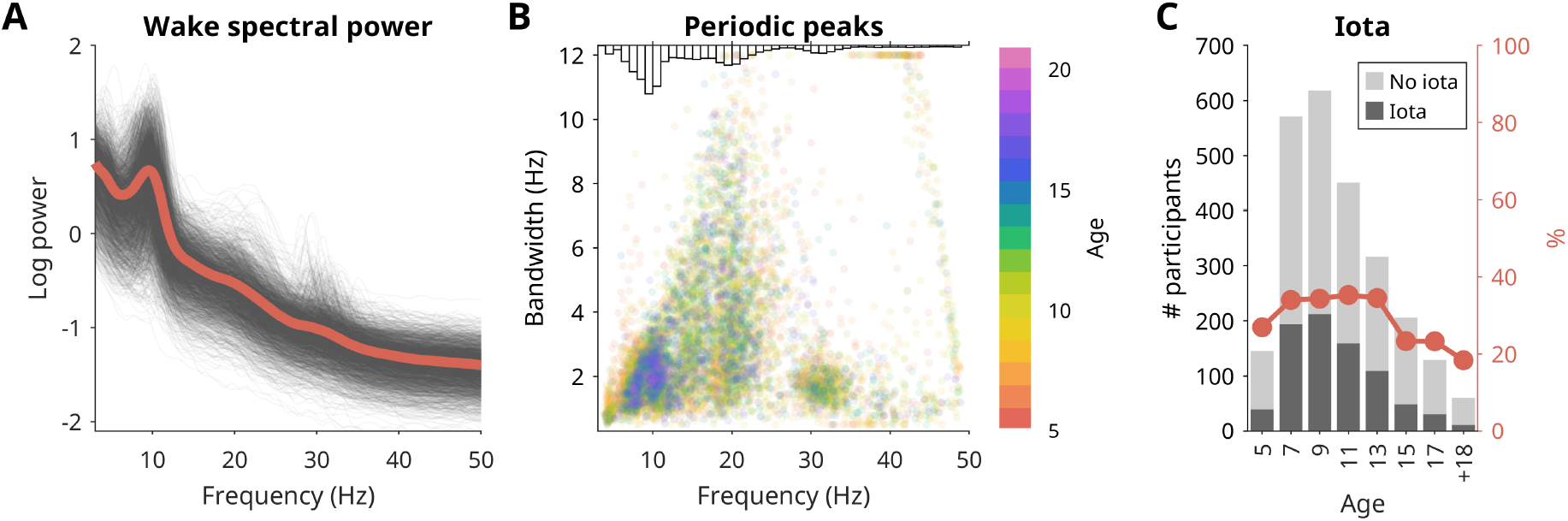
Iota in the wake HBN dataset (n=2496). **A)** All participants’ channel- and epoch-averaged log-power (black) and the average power of all participants (red). **B)** Periodic peaks detected with FOOOF from the spectra in A. Each dot represents a peak, with the x-axis indicating the center frequency, the y-axis indicating the bandwidth of the peak, and color the age of the participant. Each participant could have multiple peaks. The histogram on top indicates the total amount of peaks for each frequency. **C)** Proportion of participants where iota was identified (peak between 25-35 Hz, band-width between 0.5-4 Hz). The gray stacked bar plot indicates the absolute number of recordings with and without iota, and the red line indicates the percentage of recordings for each age bin where iota was identified.

Figure 2A provides the power spectra of all individuals. Note how periodic activity at 30 Hz is visible in some recordings but does not substantially affect the group average. Figure 2B plots the entire distribution of periodic peaks detected in the dataset, revealing the categorical separation of different oscillations. Peaks between 6 and 12 Hz with bandwidths below 4 Hz were the most prevalent, reflecting theta and alpha activity. Peaks between 12 and 25 Hz formed a more widespread cluster in both frequency and especially bandwidth, corresponding to beta.^12–14^ Then, the isolated iota cluster emerged between 25 and 35 Hz, with bandwidths generally less than 3 Hz. No substantial gamma activity was observed (>35 Hz), but this may be typical for resting-state data, as gamma is most often reported in task-based designs. Furthermore, prior studies investigating gamma report center frequencies of around 60 Hz,^15–17^ outside the investigated range. Finally, gamma is generally challenging to detect using surface EEG, and most studies investigate gamma with magnetoencephalography (MEG) or intracortical recordings.^15^ To summarize, the iota cluster emerges spontaneously in the 3-50 Hz range, separated from beta activity, and without further higher frequency activity present in the recording.

Both the beta activity observed in Figure 2B and the gamma activity reported in the literature display wide bandwidths, in the case of gamma extending up to 20 Hz.^14–17^ The 1-3 Hz bandwidths for iota oscillations are therefore quite unusual for high-frequency activity. The narrow bandwidth is also noteworthy because it explains how iota can go undetected in group averages: with center frequencies varying across a 10 Hz span, and bandwidths typically around 2 Hz, there is little overlap across individuals. Finally, the narrow bandwidth is also significant because it excludes the possibility that iota is a muscle artifact, as these have bandwidths of at least 20 Hz.^15,18^

Figure 2C plots the proportion of individuals by age with a peak between 25 and 35 Hz, and bandwidths below 4 Hz. Iota is most frequent in individuals under 14 years old (34%, 713/2101), and decreases in prevalence with age (14-18: 23%; 18-22: 18%; 89/395). The higher proportion of iota in children compared to adolescents and adults was statistically significant (2-tailed z-score test, α=5%: z= 4.453, p<.001). Despite this shift in prevalence, there was no significant correlation between age and iota amplitude among individuals with iota (Pearson’s r = -.02, p = .549), nor between age and iota frequency (r = .01, p = .831). Therefore, this suggests that iota disappears with age, rather than falling out of the defined ranges.

#### 4.1.2 Topography

Figure 3 provides the topographic distribution of iota in resting wake EEG.

**Figure 3:**
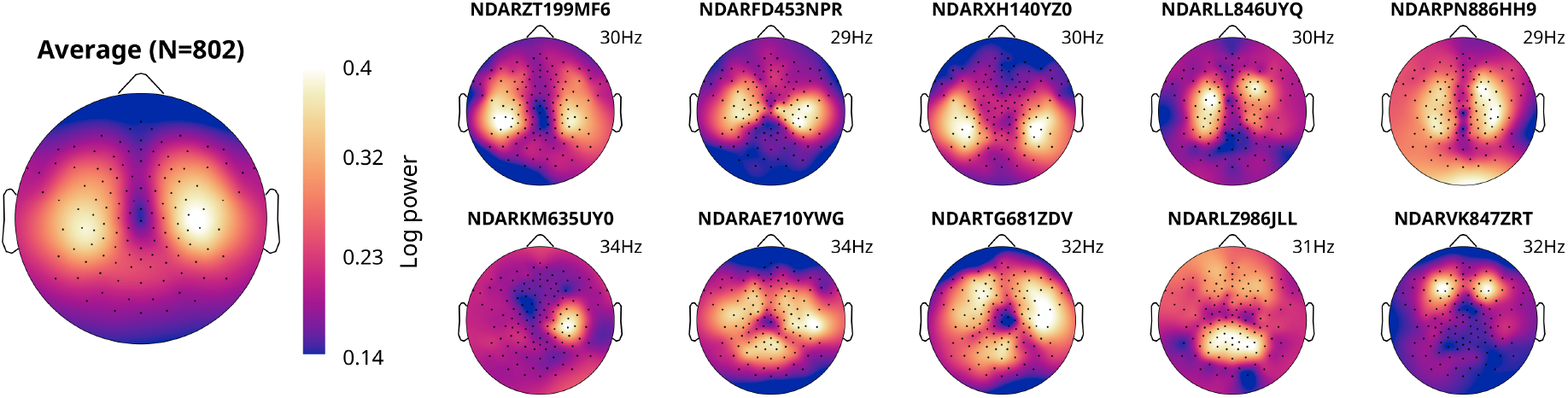
Iota topographies. Each topography provides a schematic view from above of the EEG (nose on top). Iota was periodic power averaged in custom frequency ranges (*center frequency* ± *Bandwidth/2*) for each recording. The left-most plot is the average power from all recordings where iota was detected. The smaller plots are individual examples, representing the variations in topography. The color scales of the smaller plots are individualized to maximize visibility.

Most iota activity came from temporal-parietal channels 102 and 46, corresponding to locations TP7 and TP8 in the 10-20 system. However, there were variants. Many participants had one peak stronger than the other, to the point where a few individuals only had either the left or right source of iota. Many others had a third peak in occipital areas, and a few only had this third occipital peak. Finally, some individuals had sources in bilateral frontal channels. These variants may reflect categorically distinct oscillations that by chance occupy the same band, or different mental states during the uncontrolled resting recording. Alternatively, they may simply result from interindividual differences in cortical structure. Similar topographic heterogeneity has been observed for slow sigma activity during NREM sleep,^19^ but also reflects the landscape of alpha oscillations which can originate from different cortical structures in different contexts.^20–22^

To illustrate the distinctiveness of iota, Figure 4 provides the topographies for all standard frequency bands, first as average log-power (A), then periodic power without the aperiodic component (B).

**Figure 4:**
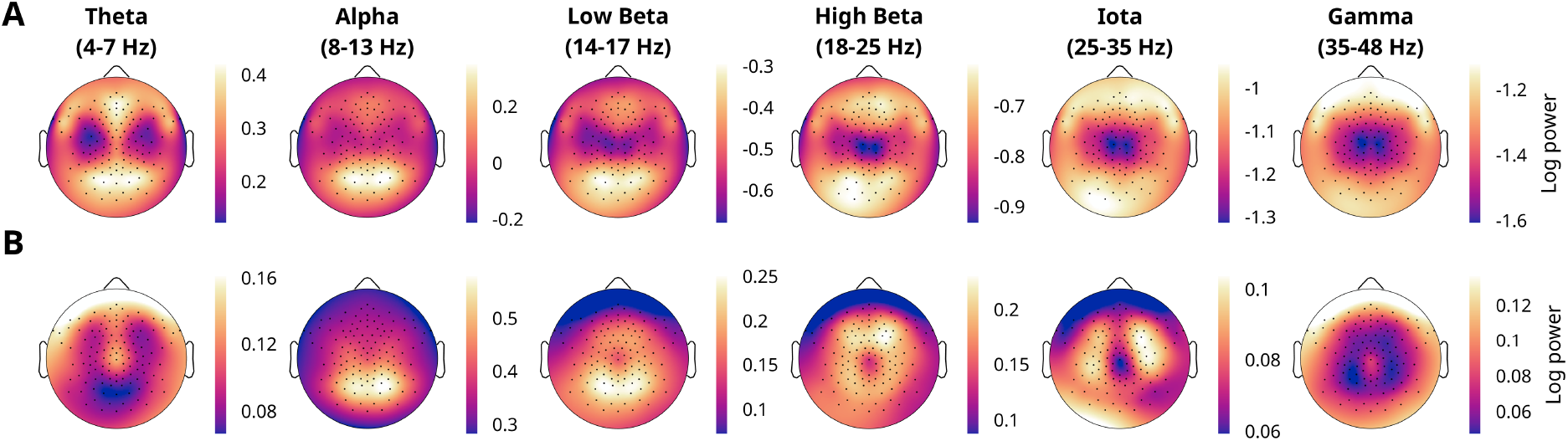
All frequency bands’ topographies. 1 Hz gaps were placed between theta, alpha, and low beta due to age-related drifts in the center frequency of alpha.^23^ **A)** Average log-power. **B)** Average periodic power (after the aperiodic signal was removed). Note how theta, high beta, iota, and gamma have different topographies from A to B.

No other frequency band originates from the same bilateral temporal-parietal channels as iota. However, high beta had bilateral frontal sources like some iota variants (Figure 3, Figure 8C).

#### 4.1.3 Waveform

To identify the waveform of iota, the preprocessed EEG traces were visually examined. Figure 5 provides example data from the participant with the highest amplitude iota peak in the HBN dataset, although the findings are consistent in other individuals examined.

**Figure 5:**
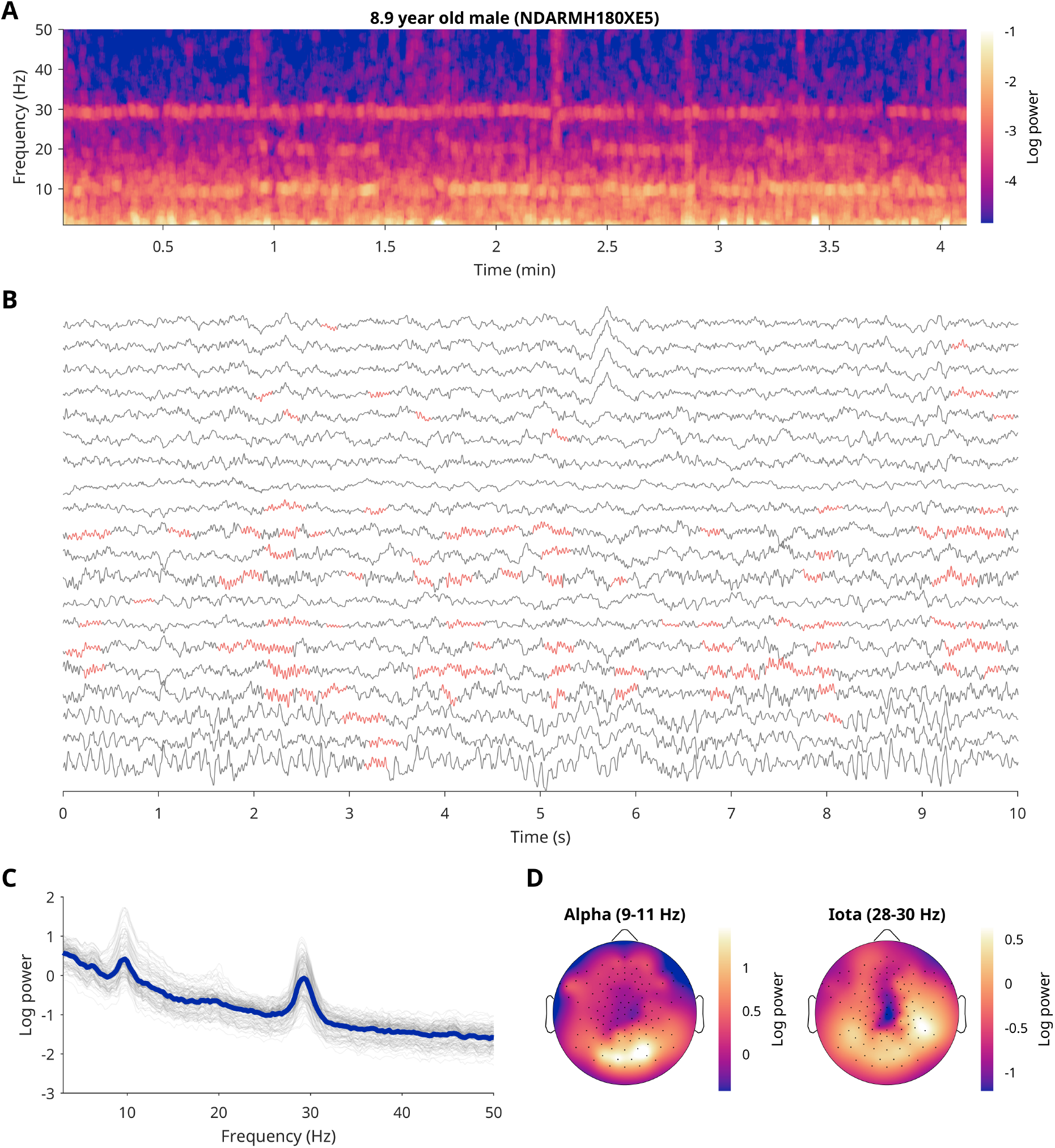
Iota in one example participant during wake. All figures originate from the same resting wake EEG recording of an 8.9-year-old male, who had the highest amplitude iota peak of the HBN dataset. **A)** Time-frequency analysis of the channel with the highest-amplitude iota, derived from a multitaper analysis using the example scripts in Cohen (2014);^25^ window length = 2 s; sampling rate = 0.2 s. **B)** 10 s example of the EEG data, with 10-20 channels (Fp1, Fp2, Fz, etc.), ordered top to bottom from front to back. Cycle-by-cycle analysis ^26,27^ was used to highlight iota bursts (in red) between 28 and 30 Hz (MonotonicityInAmplitude=.9; AmplitudeConsistency=.3; isProminent=true; MinCycles=4; ShapeConsistency=.4; FlankConsistency=.4). **C)** Log-power for all channels (gray) and their average (blue). **D)** Topographies of average log-power around the individual alpha and iota peaks.

The time-frequency plot of Figure 5A shows how iota activity is present throughout the entire recording, with minor jitter in frequency and amplitude typical of a physiological signal. Alpha is also present, and there are occasional periods with beta. Given that the participant alternated between having eyes open and eyes closed, this indicates that iota was present in both conditions.

Figure 5B provides a 10 s example of the EEG containing iota. While less prominent than alpha, it is still possible to see bursts of iota oscillations occurring in different channels at different times. This excludes the possibility that iota is merely a harmonic of a lower frequency oscillation.^24^ Note, this individual had an exceptional iota spectral peak, and yet the oscillations are still relatively hard to see in the time domain.

The fact that iota occurs in sustained bursts explains its narrow bandwidth in the power spectrum. Spectral power is measured by applying the Fourier transform to the time-domain signal, decomposing it into sine waves that can collectively reconstruct the signal. The more the signal resembles a simple sustained sine wave, the fewer sine waves are required to reconstruct it, resulting in a narrower bandwidth around the center frequency in the spectral domain. Both beta and gamma bursts have been found to appear irregularly, lasting only 1-3 cycles,^12,13,28–30^ thus producing more broadband peaks in the power spectrum.^14–17^ In other words, beta and gamma activity result in broadband spectral peaks because they are composed of short arrhythmic events, whereas the sustained oscillations of iota result in narrowband spectral peaks comparable to theta and alpha oscillations.

#### 4.1.4 Excluding artefacts

To further demonstrate that iota is not a harmonic, the iota center frequency was correlated with the alpha center frequency (Figure 6A).^24^ Pearson’s r was low (ρ=.06) and non-significant (p=.126, N=763). To demonstrate that iota was not an artefact introduced by the preprocessing, the same analysis as in Figure 2B was reproduced on data prior to any preprocessing. Iota is still detectable although less prominent (Figure 6B), underscoring how sensitive this range is to artefacts.

**Figure 6:**
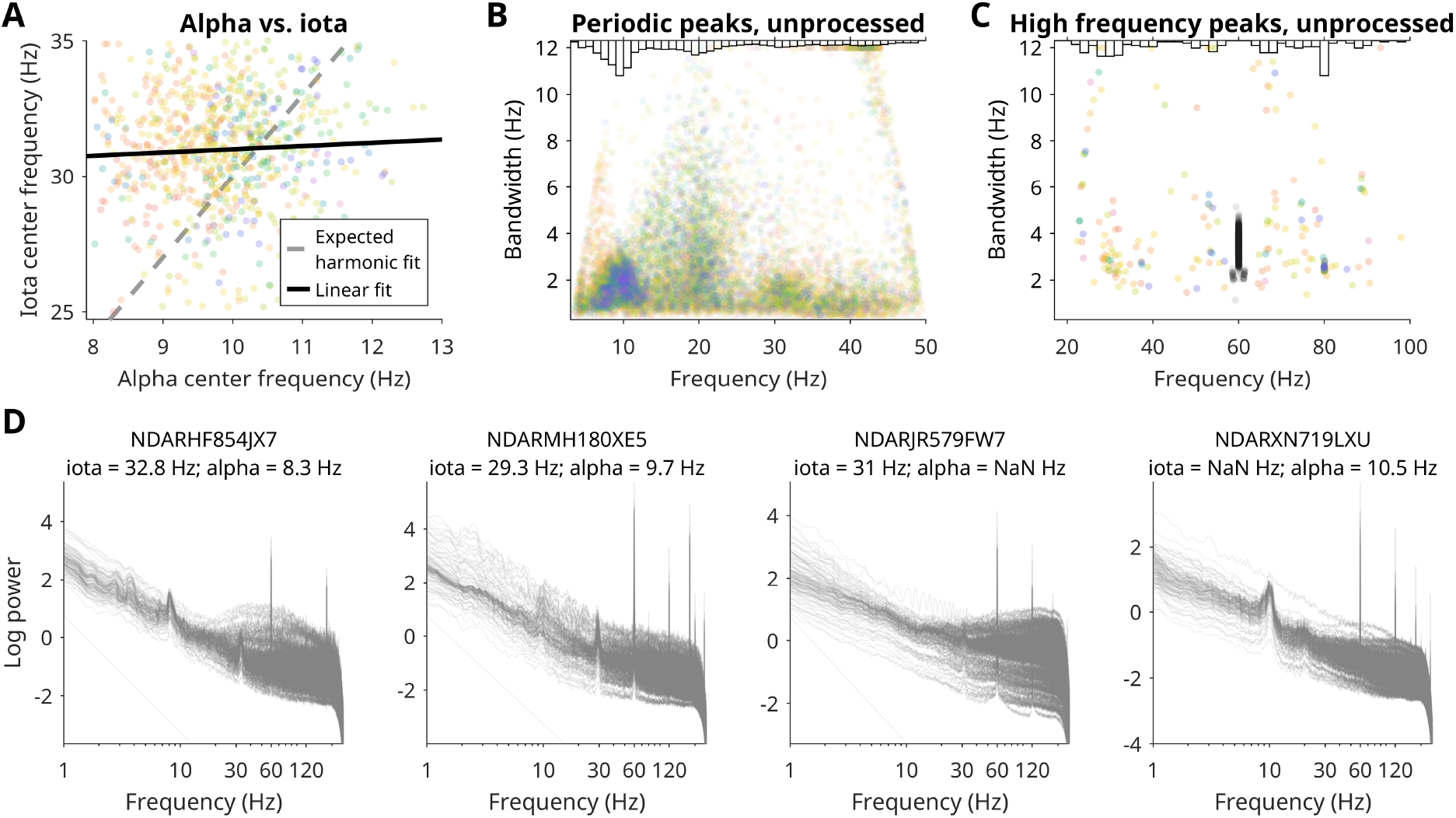
Excluding iota as an artefact. **A)** Correlations between alpha and iota. Each dot represents the periodic peaks from the average power of a single participant, color coded by age as in Figure 2B, with reds indicating the youngest children. The thick line indicates the linear fit from the data, the dotted line indicates the linear fit expected if iota had been exactly 3 × alpha, and thus a harmonic. **B)** Periodic peaks detected in data before any filtering, down-sampling, or ICA removal was applied. Artifactual epochs/channels were excluded based on aperiodic values outside of physiological ranges (see methods section 3.2.1). Then, the same procedure as for Figure 2B was applied. **C)** Periodic peaks detected in unprocessed data between 20 and 100 Hz. The plot is the same as B, except that peaks detected between 58 and 62 Hz were colored in black and excluded from the histogram for improved visibility. **D)** Untransformed power spectra of example participants, displayed on log-log scales. Unlike in B,C, no data was removed. The title indicates the participant, with their individual iota and alpha center frequencies as detected in the cleaned data.

Figure 6C applies the aperiodic fitting algorithm with a 20-100 Hz range to the unprocessed data to compare it with other known artefacts. Line noise is the most striking feature in this frequency range, appearing with extremely narrow center frequency variability, typical of artificial sources of noise. An additional artefact at exactly 80 Hz was present in some of the recordings, with minimal variation in both bandwidth and frequency. The distinctiveness of these external artefacts highlights the physiological nature of iota peaks, which instead resemble the theta-alpha peaks between 6 and 12 Hz. Other than line noise and the 80 Hz artefact, iota is the most prevalent periodic peak in this high-frequency range. Peaks detected between 35 and 40 Hz may or may not be manifestations of iota, but they are substantially less prevalent, especially after preprocessing (Figure 2B). The right-most peaks spanning across the entire bandwidth range in Figure 2B and Figure 6B, and the left-most peaks in Figure 6C, reflect aperiodic fitting imperfections due to a gradual bend in the aperiodic signal at ∼50 Hz (visible in Figure 1 and in Figure 6D).

Finally, Figure 6D provides four examples of EEG power in unprocessed data. Iota had been detected in the cleaned data of the first three, but not the last. Correspondingly, iota peaks are only visible in the first three plots.

Altogether, Figure 6 demonstrates how iota does not resemble electrical artefacts like line noise, it was not introduced by the preprocessing, and is unrelated to alpha activity.

### 4.2 REM iota

To determine whether iota depended on states of vigilance, sleep data from young healthy adults was analyzed. As an initial analysis, periodic peaks were detected in power averaged from all channels and all epochs, separately for each sleep stage (Table 1). Like this, 4 out of 19 participants were found to have iota during REM sleep, and only 1 participant had iota per remaining stage. However, closer inspection of the recordings revealed that the two “iota” peaks during NREM were harmonics of sigma (twice the frequency, same topography).

**Table 1:** Count of periodic peaks detected in sleep data from 19 young healthy adults. For each bandwidth, a peak was counted if one was detected by the FOOOF algorithm with a bandwidth below 4 Hz.

|  | W | R | N1 | N2 | N3 |
| --- | --- | --- | --- | --- | --- |
| 4-8 Hz | 8 | 6 | 8 | 11 | 13 |
| 8-13 Hz | 17 | 13 | 16 | 17 | 17 |
| 13-17 Hz | 2 | 1 | 5 | 14 | 14 |
| 17-25 Hz | 4 | 0 | 2 | 0 | 0 |
| 25-35 Hz | 1 | 4 | 1 | 1 | 1 |

In a second analysis, periodic peaks were detected for each channel and each epoch, selecting only the highest amplitude peak of each datapoint to exclude harmonics. Like this, 7 recordings had clusters during REM sleep in the iota range (Figure 7). The fact that almost twice as many participants were found to have iota in this way suggests that the previous method was not sufficiently sensitive. With one exception, all iota peaks were above 32 Hz, and an additional 3 recordings had peaks between 35 and 38 Hz, suggesting iota may be at higher frequencies during REM sleep.

**Figure 7:**
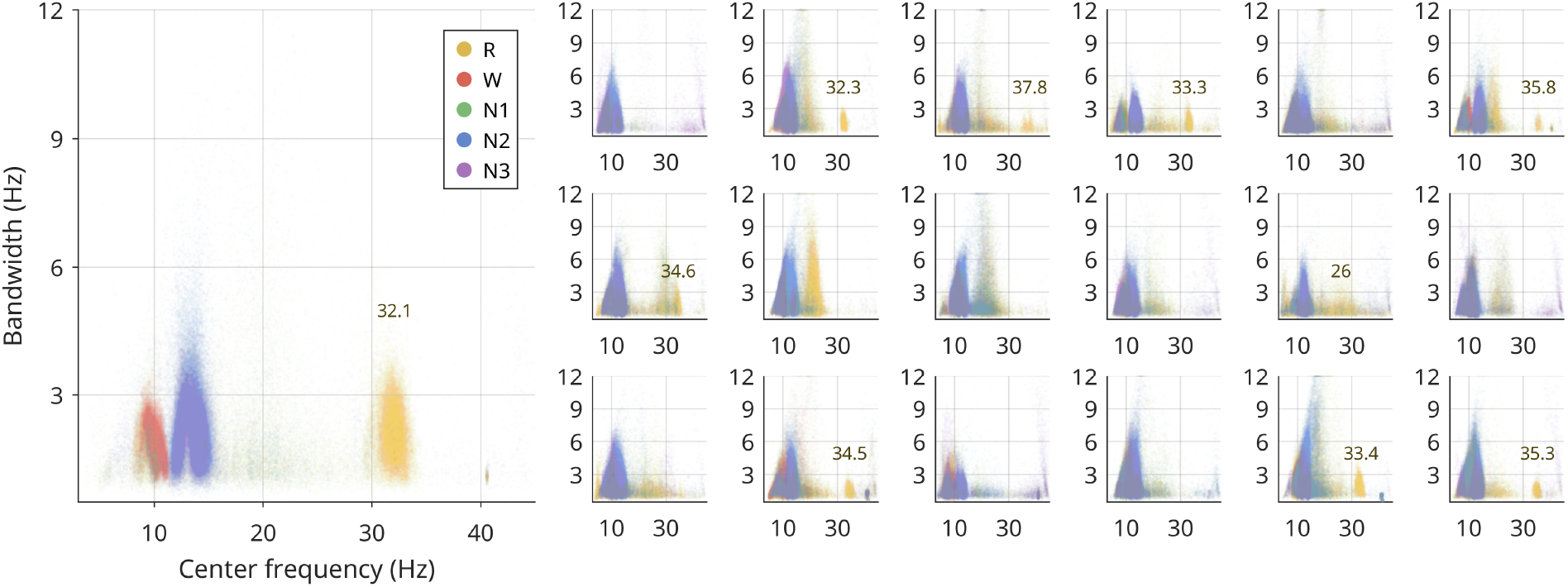
Periodic peaks during sleep in young healthy adults. Left: participant with the largest iota peak, found in REM sleep. Right: all other participants. Each plot represents all the periodic peaks from all epochs and all channels (excluding edge channels) from each participant, color coded by sleep stage. REM sleep is in yellow, wake in red, and substages of NREM sleep are in green (N1), blue (N2), and purple (N3). For each channel/epoch, only the largest peak was included. The data was plotted by stacking sleep stages, such that if there were any iota peaks detected in wake or NREM sleep, these would have appeared on top of the iota peaks in REM sleep. The iota peak frequency (text in plot) was identified by using MATLAB’s *findpeaks* on the distribution of center frequencies of peaks recorded during REM sleep between 25 and 40 Hz, for bandwidths between 0.5 and 2 Hz.

Figure 8A provides the timecourse of periodic power for the participant with the highest-amplitude iota during sleep. In this recording, iota was as characteristic of REM sleep as sigma was for NREM sleep and alpha for wake. No lower-frequency oscillation of comparable magnitude was present in this participant during REM sleep, again excluding that it could be a harmonic. Iota during REM sleep originated in bilateral frontal channels (Figure 8C). A similar topography could be found during wake, but at much lower amplitudes.

**Figure 8:**
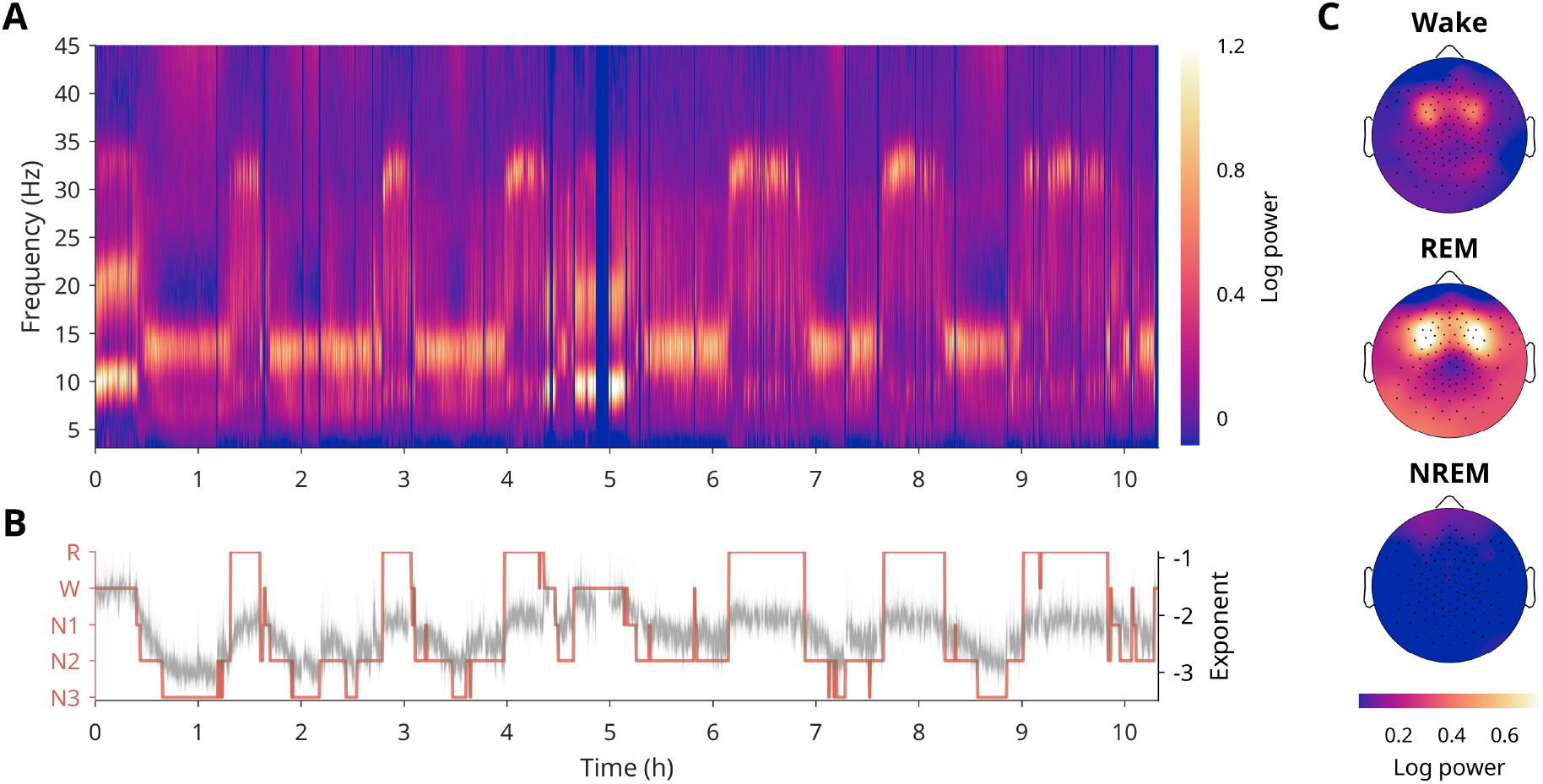
Timecourse of iota during sleep. All data from this figure originates from the same participant, who had the highest amplitude iota in REM sleep from the Zurich dataset. Other participants with iota showed similar timecourses, however iota was less prominent. **A)** Time-frequency plot of periodic power across the entire night, averaging channels. Dark vertical stripes reflect data that was removed due to artifacts. **B)** In red is the hypnogram of the sleep stages for that night based on manual scoring. In gray is the exponent of the aperiodic signal. Note, slow wave activity that characterizes NREM sleep is entirely captured by the aperiodic signal, which is why it does not appear in the time-frequency plot. **C)** Topographies of individualized iota (30-34 Hz) during wake, REM sleep, and NREM sleep.

Figure 9 provides an example of EEG data with iota oscillations during REM sleep from the participant of Figure 8. Like in wake (Figure 5B), iota occurred in sustained bursts. Iota was most prominent during periods with eye movements (i.e. phasic REM sleep), however, as can be seen from the example provided, iota also occurred in the absence of ocular activity, thus excluding the possibility that it was an eye artifact. The fact that iota occurs during REM sleep at all further excludes it from being a muscle artefact, as REM sleep is characterized by muscle atonia.^3^ Muscle twitches still occur during REM sleep but are relatively infrequent (<1/min).^31^ Furthermore, both eye movements and muscle activity are universal in healthy young adults, and should have therefore appeared in all participants.

**Figure 9:**
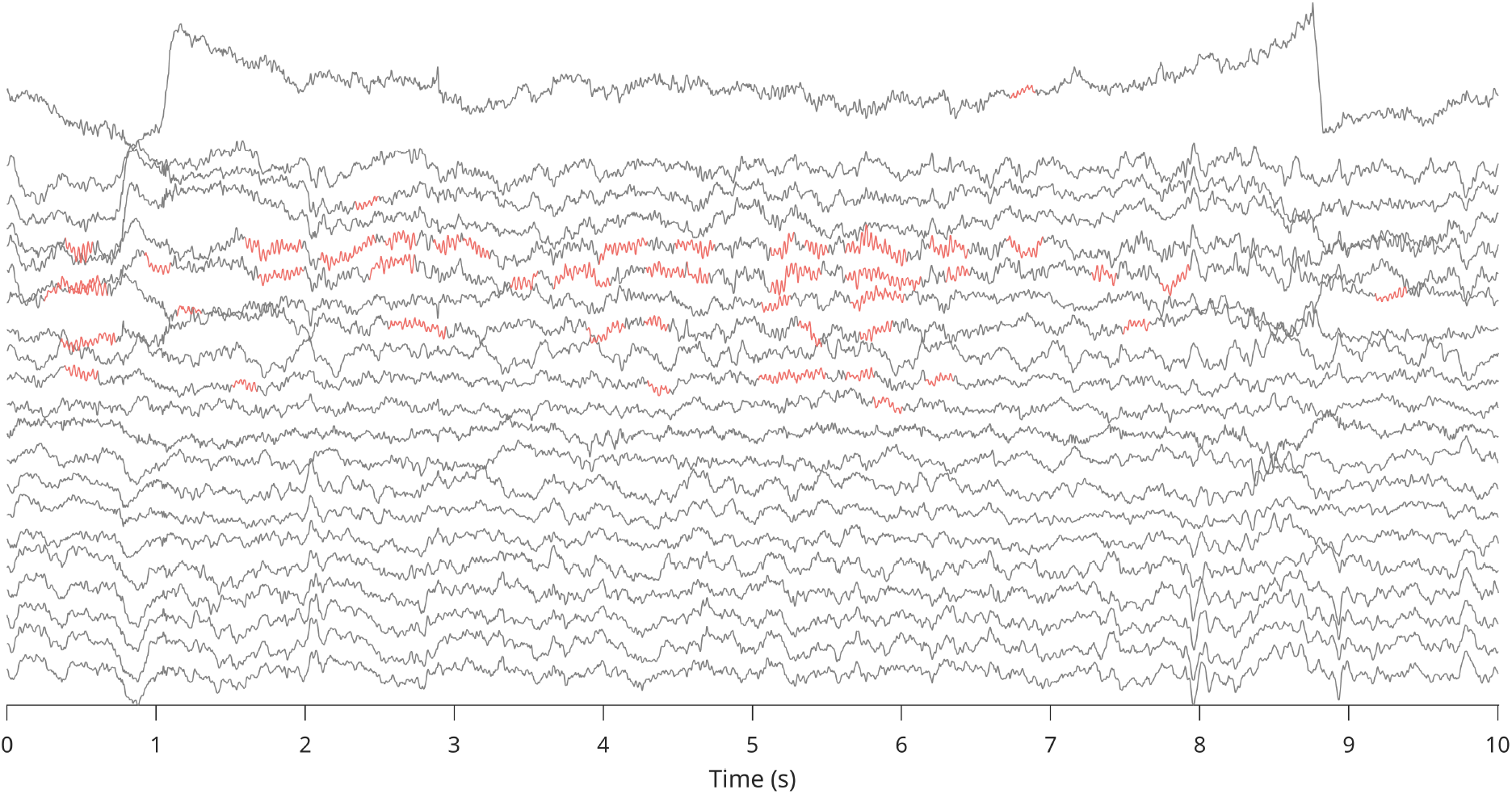
Example of iota during REM sleep. Taken from the participant of Figure 8. Channels from top to bottom correspond to standard 10-20 channels from front to back. The first channel is an EOG channel (ch 125-32). Like in Figure 5B, cycle-by-cycle analysis was used to highlight iota bursts (in red) between 30 and 34 Hz.

## 5 DISCUSSION

This paper presents evidence for distinct oscillations between 25 and 35 Hz. Iota oscillations separate themselves from other high-frequency activity by having a narrow spectral bandwidth (Figure 2B), driven by sustained oscillatory bursts (Figure 5B, Figure 9). They have a characteristic topography, albeit with a few variations (Figure 3, Figure 8C). They are only found in a minority of individuals, decreasing in prevalence with age (Figure 2C). They are state-dependent, occurring during wake and REM sleep but not NREM sleep (Figure 7, Figure 8). Given their high frequency and low amplitude, until now they have most likely been dismissed as noise. However, given their independence from both external artefacts and other EEG oscillations, their physiological variability within and across individuals, and their sleep-state dependence, they display all the typical characteristics of physiological brain oscillations.

Most of the literature into high-frequency spectral power separates beta and gamma exactly at 30 Hz, thus splitting iota in half. However, some studies have analyzed “low gamma” or “gamma1” at 25-35 Hz. Unfortunately, the aperiodic signal has an outsized effect in this range (as seen comparing Figure 4A vs. Figure 4B), and when studies find comparable results in low gamma as in neighboring bands, their results are most likely capturing the aperiodic signal.^32,33^

Distinct 30 Hz activity has been observed in infants,^34,35^ and especially young children with fragile X syndrome.^36^ However, like beta, this 30 Hz activity has a large spectral bandwidth, and when analyzed with lagged coherence it did not last more than 2-3 cycles.^35^ This suggests that iota and 30 Hz power in infants are not the same. However, it is still possible that this 30 Hz oscillation increases its frequency precision (and thus narrows its bandwidth) with development. Longitudinal data from infancy to childhood is needed to resolve this issue.

Simor et al.^37^ report effects in the 25-35 Hz range which are more likely to reflect the iota oscillations outlined here. They found that spectral power in this range was higher in phasic REM sleep (with eye movements) compared to tonic REM sleep (without eye movements) in 4–8-year-old children, but this effect was absent in young adults. This could therefore reflect a greater incidence of iota in phasic REM sleep, at least in children who are more likely to have iota in the first place. Other papers may show evidence of iota in their spectrograms, but without further comment in the text.^38^

Substantially more research is needed to understand iota oscillations. Unlike lower frequencies, however, iota is challenging to detect. The fact that it was not found in most individuals could be due to it not being a universal oscillation, or because it is often too low-amplitude to be visible in the surface EEG, or it is not always present during a 5 minute resting wake recording. Even in participants with the highest-amplitude iota, it is difficult to see in the time domain and is therefore best identified in the spectral domain. Importantly, iota should be detected by first removing the background aperiodic signal. Iota will likewise be completely masked if there is excessive muscle activity or other high-frequency artefacts. Analyses also need to control for manifestations in the iota frequency range that may merely be harmonics of non-sinusoidal low-frequency oscillations.^24^ Furthermore, analyses should differentiate between participants who do or do not have iota, and would benefit from adapting the frequency range to each individual.

Most questions related to iota remain unanswered. The fact that the HBN dataset is a highly heterogeneous patient population with uneven age distribution limits the generalizability of these results. Likewise, the sleep dataset analyzed here was small and limited to young adults. More thorough investigations with appropriate datasets are needed. Important open questions also include: what is the functional role of iota oscillations? How do they relate to behavior? Why do only some individuals have them, why do they decrease in prevalence with age, and how do they relate to pathology? Do they exist in other species? What are the neural sources generating iota, and why do they change across individuals and possibly states of vigilance? Other than wake and REM sleep, in which conditions are they more likely to appear? Now that iota oscillations have been identified, future research can begin to answer these questions.

## 6 ACKNOWLEDGEMENTS

A special thank you goes to Simon Accascina, Maria Dimitriades, Thomas Andrillon, and Valeria Jaramillo for proof-reading and feedback on the manuscript. Thank you also to Prof. Reto Huber and the Children’s Hospital of Zurich for hosting me during this project. For the Zurich sleep dataset, Elias Meier helped collect the data, Selina Schuele scored it, and Hans-Peter Landolt hosted the experiments at the University of Zurich. Data collection of the Zurich dataset was funded by the SleepLoop Flagship project of Hochschulmedizin Zürich, with additional funding from the Swiss National Science Foundation (320030_179443) and Hirnstiftung. Thank you also to the Child Mind Institute for making their data freely available.

*For any questions or comments, please feel free to contact me at publishing@sophia*.*science. If you know of any relevant literature on 30 Hz activity, please send it my way*.

